# Short-term statin treatment reduces, and long-term statin treatment abolishes chronic vascular injury by radiation therapy

**DOI:** 10.1101/2023.09.20.558723

**Authors:** Karima Ait-Aissa, Xutong Guo, Madelyn Klemmensen, Linette N. Leng, Olha M. Koval, Isabella M. Grumbach

## Abstract

**Background:** The incidental use of statins during radiation therapy has been associated with a reduced long-term risk of developing atherosclerotic cardiovascular disease.

**Objectives:** Determine if irradiation causes chronic vascular injury and whether short-term administration of statins during and after irradiation is sufficient to prevent chronic injury compared to long-term administration.

**Methods:** C57Bl/6 mice were pretreated with pravastatin for 72 hours and then exposed to 12 Gy x-ray head-and-neck irradiation. Subsequently, they received pravastatin either for one additional day or for one year. Carotid arteries were tested for vascular reactivity and altered gene expression one year after irradiation.

**Results:** Treatment with pravastatin for 24 hours reduced the loss of endothelium-dependent vasorelaxation and protected against enhanced vasoconstriction after IR. It reduced the expression of some markers associated with inflammation and oxidative stress and modulated that of subunits of the voltage and Ca^2+^ activated K^+^ (BK) channel in the carotid artery one year after irradiation. Treatment with pravastatin for one year completely reversed the changes caused by irradiation.

**Conclusions:** In mice, short-term administration of pravastatin is sufficient to reduce chronic vascular injury after irradiation. Long-term administration eliminates the effects of irradiation. These findings suggest that a prospective treatment strategy involving statins could be effective in patients undergoing radiation therapy. The optimal duration of treatment in humans has yet to be determined.

## INTRODUCTION

Radiation therapy has been associated with an elevated risk of developing vascular disease ^1^. Following radiation therapy for head-and-neck cancer, the risk for carotid stenosis is strongly increased ^2-5^ because the carotid artery is adjacent to the targeted lymphatic structures and usually included in the radiation field ^2-5^. In patients who had undergone radiation therapy for head-and-neck cancer, the rate of progression of carotid artery stenosis to >50% was 3-fold higher than in patients who had not received radiation therapy but were matched for the severity of carotid artery stenosis at baseline ^6^.

Although, to date, no drugs have been designed specifically to suppress IR-induced vascular disease, the incidental use of statins at the time of, or after, radiation therapy for head-and-neck cancer has been associated with a lower risk of stroke ^7,8^. In spite of these promising observations, prospective clinical trials directly testing whether statin treatment reduces vascular disease in this population have not been undertaken, in part because of the long interval between radiation therapy and clinical events ^7^. Given these considerations, studies in preclinical models can provide valuable information. We recently reported that in C57Bl/6 mice, treatment with pravastatin for up to 10 days after irradiation (IR) prevented endothelial dysfunction, mitochondrial damage, and endothelial senescence ^9^. In the current study, we sought to determine whether IR induces chronic injury by upregulating mediators of inflammation and oxidative stress, as reported in human subjects ^10^, and whether statin therapy protects from chronic vascular injury. Additionally, we tested two regimens of pravastatin administration to determine the duration of therapy necessary to prevent vascular injury. Specifically, we looked at a one-day treatment after IR (short-term) and continuous one-year treatment after IR (chronic, long-term). We examined the effects on dilation and constriction of the common carotid artery, and the expression of markers of inflammatory and oxidative stress, and of structural components of the vascular wall.

## MATERIALS and METHODS

### Mice

All experimental procedures were approved by the Institutional Animal Care and Use Committees of both the University of Iowa and the Iowa City VA Health Care System and complied using the standards of the Institute of Laboratory Animal Resources, National Academy of Sciences. Male and female C57BL/6J mice were obtained from Jackson Laboratories (#000664). Mice were housed at 23°C on 12:12-hour light/dark cycle. All mice were between 12-16 weeks of age at the time of treatment.

#### Statin treatment

The mice were divided into 3 groups: 1) the control group received vehicle only (filtered tap water); 2) the short-term treatment group received pravastatin in drinking water starting 72 hours before IR and for additional 24 hours after IR; and 3) the long-term treatment group received pravastatin in drinking water starting 72 hours before IR and through 12 months after IR. Pravastatin was given orally in drinking water, which was provided ad libitum, resulting in a dose of approximately 30 mg/kg/day ^11^.

#### Radiation therapy of mice

Mice were anesthetized with isoflurane and irradiated using the XStrahl Small Animal Radiation Research Platform (SARRP) with a single anterior-posterior beam and a beam quality of 0.67 mm Cu, as previously described ^9^. Sham-treated mice were anesthetized but did not undergo IR.

The dose rate was 3.6 Gy/min and calibrated at 2 cm depth in water, in accordance with the AAPM TG-61 protocol. A dose of 12 Gy X-ray, which equates to EQD2 dose of 36 Gy (α/β of 3), was delivered to the head and neck in a single session. Simulation was performed by computed tomography. The accuracy of dosimetry by the SARRP was ensured by quarterly measurements of the ion chamber by a medical physicist. ^12^.

### Vascular reactivity

Arterial rings were prepared from the common carotid and second-branch mesenteric resistance arteries. Their isometric tension was measured after mounting the rings in a small vessel dual chamber myograph. Following equilibration in Krebs solution bubbled with CO_2_ at 37°C and at pH 7.4 for 30 minutes, the rings were stretched to their optimal physiological lumen diameter for one hour to develop active tension. The rings were then pre-constricted with phenylephrine (PE, 3X10^-5^ M), after which they were treated with acetylcholine (ACh, 10^-8^ - 3X10^-5^ M) or sodium nitroprusside (SNP, 10^-8^ - 3X10^-5^ M) to generate cumulative concentration-response curves. Mesenteric resistance arteries served as control arteries from a vascular bed outside the radiation field and were treated in the same fashion.

### Quantitative Real-Time PCR

Total RNA extracted from carotid artery lysates was reverse transcribed and amplified in a ViiA 7 Real-Time PCR System (Applied Biosystems, Foster City, CA). Primers were designed by Integrated DNA Technologies to amplify the following genes: NF κB-p65, TNFα, intercellular adhesion molecule 1 (ICAM1), endothelial nitric oxide synthase (eNOS), NADPH oxidases 2 and 4 (NOX2, NOX4), myosin heavy chain 11 (MYH11), myosin light-chain kinase (MLCK), smooth muscle actin (SMA), collagen type I α2 chain (COL1A2), potassium calcium-activated channel subfamily M alpha 1 (KCNMA1), potassium calcium-activated channel subfamily M regulatory beta subunit 1 (KCNMB1) and ribosomal 18S or β-actin (internal control). Primers sequences are provided in Supplemental Table 1.

### Statistical analysis

Data were expressed as mean ± standard error of the mean (SEM) and analyzed using the GraphPad Prism 9.0 software. All data sets were analyzed for normality and equal variance. Kruskal-Wallis and Dunn’s post hoc tests were used for data sets when a normal distribution could not be assumed. Comparisons were made to vehicle-irradiated conditions. One-way ANOVA, followed by Tukey’s multiple comparison test, was used for data sets with a normal distribution. Two-way ANOVA followed by Tukey’s multiple comparison test was used for grouped data sets. A p-value <0.05 was considered significant.

## RESULTS

### Short-term treatment with pravastatin partially alleviates abnormal vascular reactivity at one year following IR, while long-term treatment prevents it

At the one-year mark after IR, we measured the dilation and constriction of the common carotid artery in response to different treatments: acetylcholine (ACh) for endothelium-dependent dilation, sodium nitroprusside (SNP) for endothelium-independent dilation, and phenylephrine (PE) for vasoconstriction.

At this time point, we observed that relaxation in response to ACh was significantly reduced in the common carotid arteries of IR mice compared to non-irradiated sham control (nIR) mice (**Figure 1A, B**). In mice that had received short-term pravastatin treatment (one day), endothelium-dependent relaxation was impaired compared to nIR mice and improved compared to IR mice that had not received pravastatin. (**Figure 1A, C**). However, IR mice that underwent long-term treatment (one year) with pravastatin exhibited the same level of endothelium-dependent relaxation than mice in the nIR group, indicating complete preservation (**Figure 1A, D**). Vasoconstriction in response to phenylephrine was increased in the IR group compared to the nIR group (**Figure 1E, F**), and both short- and long-term treatment with pravastatin normalized the vasoconstriction after IR (**Figure 1E, G, H)**.

**Figure 1:**
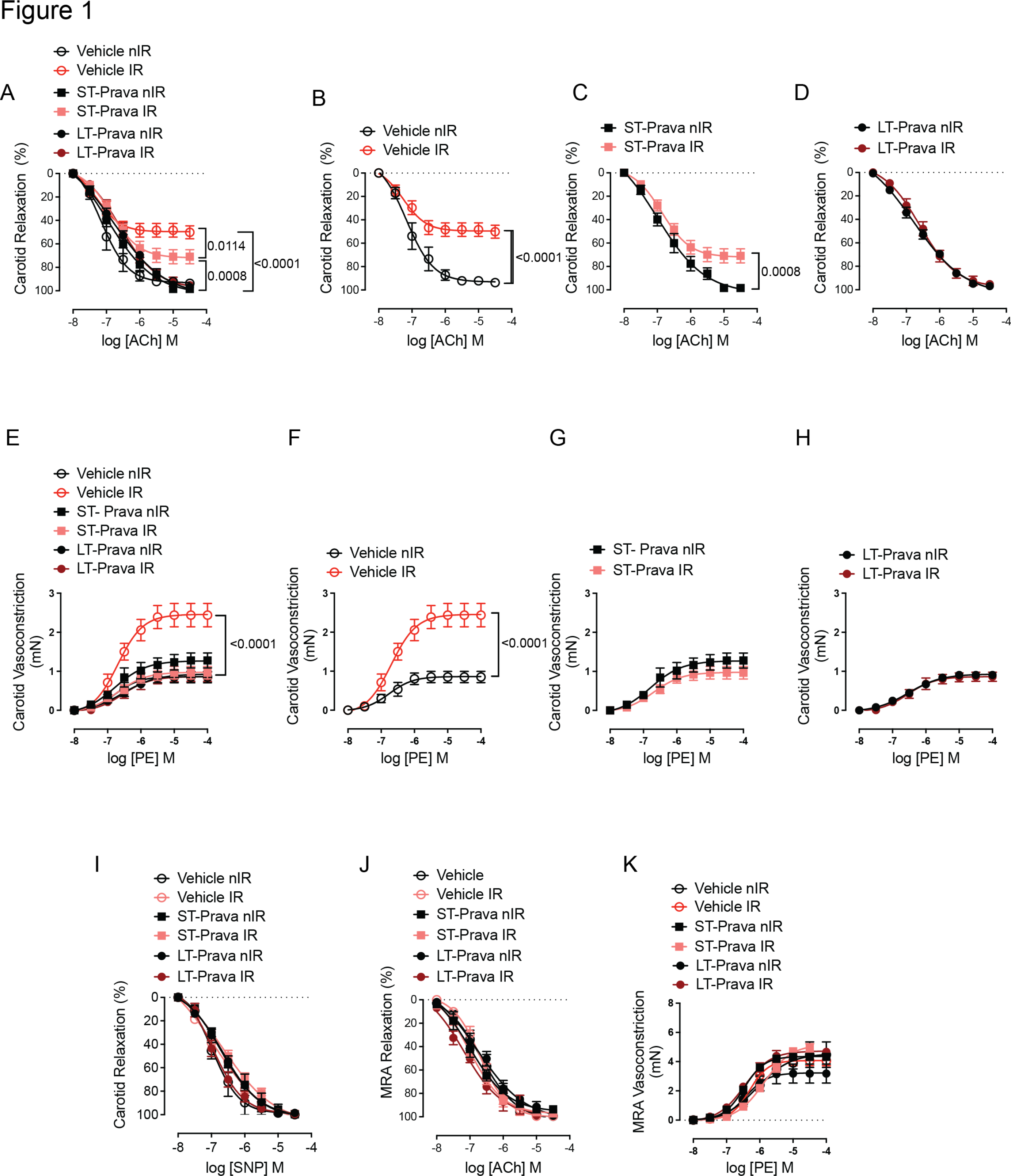
Pravastatin treatment preserves endothelial function *ex vivo* following head- and-neck IR. **(A–D)** Effects of short-term (ST, one-day) and long-term (LT, one-year) pravastatin (Prava) treatment after head-and-neck irradiation (IR, 12Gy X-ray), on endothelium-dependent relaxation of the common carotid artery in response to acetylcholine (ACh). Pravastatin treatment (30 mg/kg/day) for both groups was started at 72 hours before IR. C57BL/6J mice were subjected to **(A, B)** vehicle only treatment, **(A, C)** ST-pravastatin treatment, or **(A, D)** LT-pravastatin, with IR or with sham treatment (non-irradiated, nIR), and relaxation was tested at one year after IR. **(E-H)** Effects of **(E, F)**, vehicle, **(E, G)** ST pravastatin treatment, or **(E, H)** LT pravastatin treatment on constriction of the common carotid artery in response to phenylephrine (PE), in mice treated as in A-D, respectively. **(I)** Effects of vehicle, ST or LT pravastatin treatment on endothelium-independent relaxation of the common carotid artery in response to sodium nitroprusside (SNP). **(J)** Effects of vehicle, ST or LT pravastatin treatment on endothelium-dependent relaxation of mesenteric resistance arteries (MRAs). (**K**) Effects of vehicle, ST or LT pravastatin treatment on constriction of the common carotid artery in response to PE. n=4 mice per group for Vehicle nIR, Vehicle IR and ST-Prava nIR, 8 for ST-Prava IR, 6 for LT-Prav nIR and 7 for LT-Prava IR. p values were determined using repeated measures 2-way ANOVA followed by Tukey’s post-Hoc test.

Control experiments assessing the effects of IR and statins on endothelium-independent dilation by SNP confirmed that neither affected carotid relaxation (**Figure 1I**). Moreover, in mesenteric resistance arteries outside the radiation field, neither dilation in response to Ach nor constriction in response to PE was affected by either IR or statin treatment (**Figure 1J, K**).

### Short-term treatment with pravastatin reduces changes in the expression of markers of inflammation, oxidative stress, and structural wall remodeling at one year after IR, while long-term treatment prevents them

To identify the protective effects of short-vs long-term treatment with pravastatin, we used quantitative RT-PCR to analyze the expression of markers associated with inflammation, oxidative stress, and structural remodeling. At the one-year mark after IR, carotid arteries from mice treated with vehicle and IR exhibited higher levels of mRNAs encoding NFkB-p65, TNF-α, and ICAM1 compared to vehicle-treated nIR mice. However, these effects were abolished by long-term treatment and mildly reduced by short-term treatment with pravastatin (**Figure 2A-C**). In contrast, the mRNA for the endothelial nitric oxide synthase (eNOS), which regulates nitric oxide synthesis, was significantly lower in carotid arteries from IR mice compared to nIR mice (**Figure 2D**). Both pravastatin treatments prevented this difference, with the long-term treatment showing a more significant effect. Additionally, the expression of NOX2 and NOX4 mRNAs, which encode NADPH oxidases expressed in the vascular wall, was higher in carotid arteries from IR mice compared to nIR vehicle-treated mice (**Figure 2E, F**). Long-term pravastatin treatment blocked the effect of both NOX2 and NOX4, while short-term pravastatin treatment specifically blocked the effect of NOX4.

**Figure 2:**
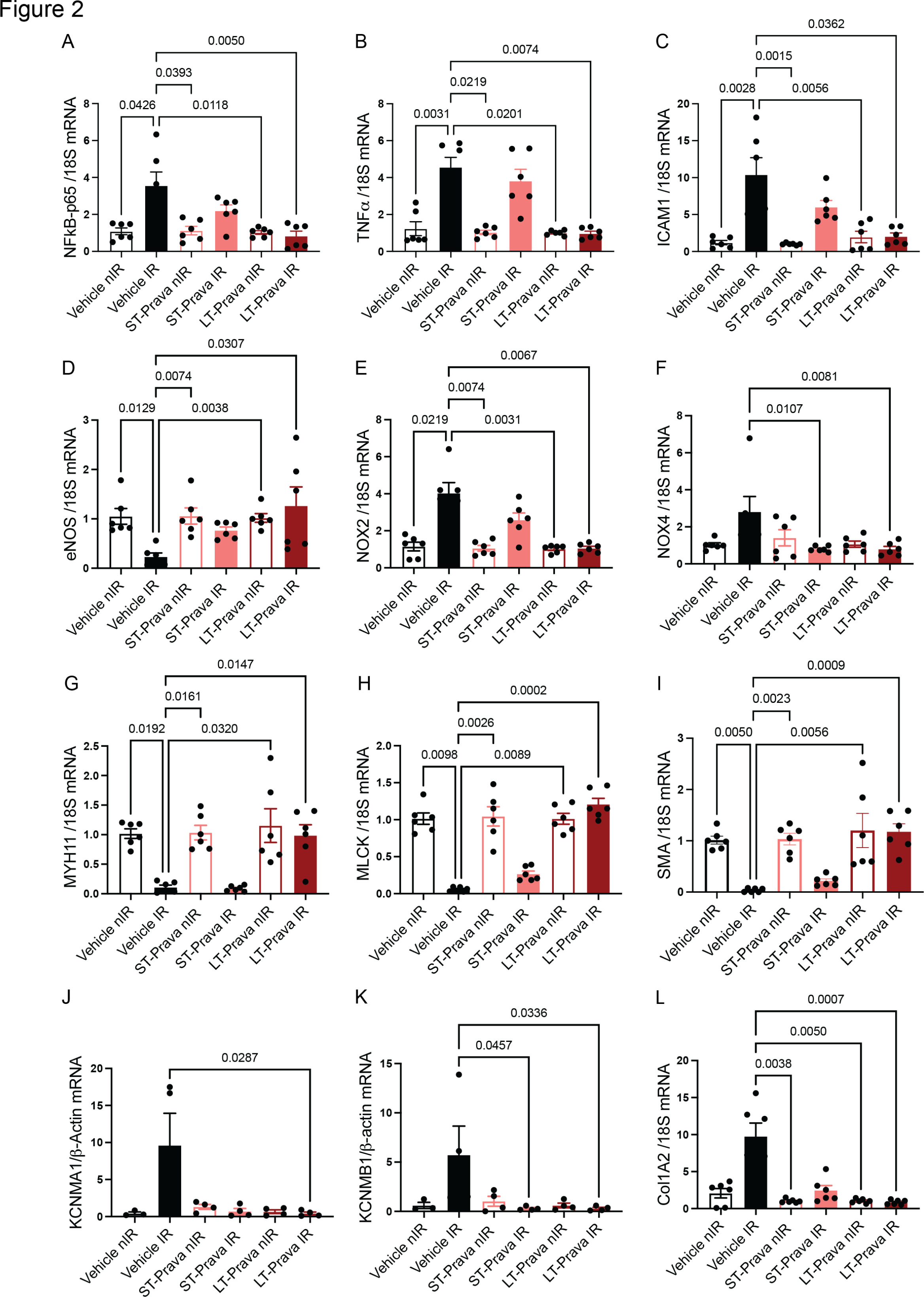
Pravastatin reduces IR-induced expression of genes in inflammatory oxidative stress and vascular wall structural pathways. All panels compare gene expression in common carotid arteries of C57BL/6J mice that were pretreated with vehicle or pravastatin for 72 hours, subjected to irradiation (12 Gy), and then underwent short-term (ST, one-day) or long-term (LT, one year) pravastatin treatment. Quantitative RT-PCR was performed at one year after IR. Quantitative RT-PCR for **(A)** NFκB-p65, **(B)** TNFα, **(C)** ICAM1, **(D)** eNOS, **(E)** NOX2, **(F)** NOX4, **(G)** myosin heavy chain 11 (MYH11), **(H)** myosin light-chain kinase (MLCK), **(I)** smooth muscle actin (SMA), **(J)** potassium calcium-activated channel subfamily M alpha 1 (KCNMA1), **(K)** potassium calcium-activated channel subfamily M regulatory beta subunit 1 (KCNMB1) and **(L)** collagen type I alpha 2 chain (COL1A2). n=7 mice per group for A-I, L, n=4 for J, K. p values were determined using the Kruskal-Wallis test.

Furthermore, IR reduced the levels of mRNAs encoding smooth muscle genes, including myosin heavy chain 11, myosin light-chain kinase and smooth muscle actin (**Figure 2G-I**). Only long-term pravastatin treatment abolished this effect. We made similar findings for the SM22α (data not shown).

Given that vasoconstriction in response to phenylephrine was increased in the IR group compared to the nIR group, and both short- and long-term treatment with pravastatin normalized it, we investigated whether the expression of subunits of the voltage and Ca^2+^ activated K^+^ (BK) channel is altered under these conditions. Previous studies suggested that altered posttranscriptional regulation and decreased mRNA levels of Ca^2+^-activated K^+^ (BK) channels in smooth muscle cells contribute to increased constriction after IR ^13-16^. Here, we detected significant increases in mRNA levels of the BK channel subunits KCNMA1 and KCNMB1 with IR. Both long- and short-term treatment with pravastatin abolished this effect (**Figure 2J, K**).

Finally, we assessed the impact of pravastatin treatment on collagen expression induced by IR. qRT-PCR showed that expression of the collagen type I α2 chain was elevated after IR in vehicle-treated mice, and pravastatin treatment reduced this effect (**Figure 2L**)

## DISCUSSION

In this study, we aimed to determine whether vascular injury resulting in changes in vascular reactivity, and expression of markers related to inflammation and oxidative stress could be detected one year after IR. Additionally, we investigated the protective effects of treatment with pravastatin, either for one day only or throughout the entire period after IR, against vascular injury. Our study yielded three significant findings. Firstly, we observed impaired endothelium-dependent dilation and increased constriction in the common carotid artery one year after IR. Secondly, there was an elevation in the expression of mRNAs encoding inflammatory markers, NOX2, and NOX4 and BK channel subunits KCNMA1 and KCNMB1, while mRNAs encoding eNOS and smooth muscle structural proteins showed lower expression in IR mice compared to nIR mice. Thirdly, long-term treatment with pravastatin throughout the post-IR period eliminated the effects of IR, while even a one-day short-term treatment normalized constriction and the expression of NOX4, KCNMA1 and KCNMB1, with some effects on endothelium-dependent dilation, and the expression of other pro-inflammatory proteins and collagen mRNA.

In patients with head-and-neck cancer, IR drives the progression of carotid arterial disease, leading to carotid artery stenosis and stroke ^17^. However, there are currently no specific therapies available to treat IR-induced cardiovascular disease. As a result, preventive treatments such as aspirin, colchicine, and statins have been proposed ^18^. Interestingly, the incidental use of statins lowered the risk of stroke in patients after IR for head-and-neck cancer^7^. However, in adult survivors of childhood cancer, atorvastatin did not improve endothelial function or arterial stiffness, possibly because the atherosclerotic risk in this population is lower than those in adult cancer patients ^19^. Studies in humans have identified changes in the expression of ICAM-1, VCAM-1, E- and P-selectin, eNOS, and increased NF-κB signaling within a span of up to 10 years after IR ^10,20,21^. Preclinical studies in mice are expected to provide additional insights since the same mechanisms of arterial inflammation and oxidative stress have been implicated in IR-related vascular disease. However, in preclinical models, altered vascular reactivity, inflammation and oxidative stress have been reported within a few weeks after treatment ^22-25^. Our experiments, conducted one year after IR, aimed to bridge the knowledge gap regarding long-term effects in preclinical models. Our findings confirm the presence of inflammation, potentially driven by NF-κB, oxidative stress, and the expression of pro-inflammatory genes at this time point, which are reduced by statin treatment. IR impairs endothelial-dependent vasodilation through NO-dependent pathways ^16,26^ in humans and in preclinical models ^27^. Endothelial dysfunction is a strong predictor of atherosclerotic risk and adverse cardiovascular events, including stroke ^28,29^. Statin therapy maintains NO production by preserving expression of the eNOS mRNA ^30^. Statin therapy also protects against uncoupling of eNOS, thereby decreasing ROS production. Our findings suggest that statin therapy by reducing the expression of NOX proteins ^31^ lowers ROS production and increases eNOS levels, resulting in preserved vasodilation after IR.

Although less is known about the effects of IR on vasoconstriction compared to vasodilation, previous studies indicate that altered posttranscriptional regulation and decreased mRNA transcription of BK channel subunits in smooth muscle cells drive increased constriction ^13-16^. Opposite to our predictions, mRNA levels of the BK channel subunits KCNMA1 and KCNMB1 were higher at one year after IR compared to control nIR conditions. Both short-term and long-term statin treatments normalized mRNA expression. The findings will warrant further studies of BK channel protein levels in smooth muscle cells of the vascular wall after IR. This will be critical because the conclusions in previous studies were solely drawn from electrophysiological studies with BK and PKC inhibitors and quantification of BK channel subunit mRNA transcripts. Moreover, to get a complete picture, there is a need to study other channels relevant for vasoconstriction, including K_v_, KATP and L-VDCC channels.

Increased fibrosis is a well-established characteristic of pathology in normal tissue after IR ^32,33^. In our studies, pravastatin abolished collagen expression, which may have contributed to the normalization of vasoreactivity by this treatment.

Our data indicate that treatment with statins during and after IR can mitigate the detrimental effects of IR on vascular function and atherosclerotic risk factors. Administering pravastatin for only one day after IR improved dilation, normalized constriction, and blocked increased expression of NOX4 and BK channel subunit KCNMB1 and attenuated changes in mRNA levels of other markers caused by IR. These findings suggest that a brief course of statin treatment may be sufficient to protect against certain aspects of vascular injury following IR. Further studies will be essential to determine the ideal duration of statin treatment and to fully establish the molecular mechanisms that contribute to enhanced constriction after IR. The knowledge gained from our study in mice can guide the development of trials involving human subjects.

## ABBREVIATIONS

ACh: acetylcholine
BK channel: Ca^2+^ activated K^+^ channel
COL1A2: collagen type I α2 chain
eNOS: endothelial nitric oxide synthase
ICAM1: intercellular adhesion molecule 1 IR irradiation, irradiated
KCNMA1: potassium calcium-activated channel subfamily M alpha 1
KCNMB1: potassium calcium-activated channel subfamily M regulatory beta subunit 1
MLCK: myosin light-chain kinase
MYH11: myosin heavy chain 11
nIR: non-irradiated
NOX2: NADPH oxidases 2
NOX4: NADPH oxidases 4
PE: phenylephrine
SE: standard error
SMA: smooth muscle actin
SNP: sodium nitroprusside
SARRP: small animal radiation research platform

## ACKNOWLEDGMENTS

We thank Dr. Christine Blaumueller of the Scientific Editing and Research Communication Core at the University of Iowa for critical reading of the manuscript and editorial assistance. Experiments in the Radiation and Free Radical Research Core Facility reported in this publication were supported by the National Cancer Institute of the National Institutes of Health under Award Number P30CA086862.

